# One hundred years of climate change in Mexico

**DOI:** 10.1101/496216

**Authors:** Angela P. Cuervo-Robayo, Carolina Ureta, Miguel A. Gómez-Albores, Anny K. Meneses-Mosquera, Oswaldo Téllez-Valdés, Enrique Martínez-Meyer

## Abstract

Spatial assessments of historical climate change are of paramount importance to focus
research and conservation efforts. Despite the fact that there are global climatic databases available at high spatial resolution, they present some shortcomings to evaluate historic trends of climate change and their impacts on biodiversity. These databases span over a single period in the late 20^th^ and early 21^th^ centuries and their quality and reliability in many regions is compromise because they have not been produce with all information available for all regions. Therefore, in this contribution we developed climatic surfaces for Mexico for three periods that cover most of the 20^th^ and early 21^th^ centuries: *t*_*1*_-1940 (1910-1949), *t*_*2*_-1970(1950-1979) and *t*_*3*_-2000 (1980-2009), and characterize climate change rates of the bio-geographic provinces of Mexico via a linear trend analysis of monthly values and a Mann-Kendall analysis. Our results indicate that rates of change and trends have not been uniform across Mexico: Nearctic provinces had suffered higher and more consistent trends of change than southern tropical regions. Central and southern provinces cooled down at the beginning of the 20^th^ century, but warmed up consistently since the 1970s. Precipitation has generally increased throughout the country, being more notorious in northern provinces. We aim to provide modellers with a set of climate surfaces that may help decision-making to improve management strategies for biodiversity conservation.

## Introduction

Climate change has been recognized as one of the major drivers of biodiversity loss in recent years [1] due to its strong effect on demographic, geographic and ecosystem processes [1,2,3,4,5]. Additionally, climate change acts synergistically with other environmental degradation factors, such as habitat loss, pollution, overexploitation, and invasive species (Saunders et al. 1991; Guisan et al. 2014). Currently, studies dealing with climate change and biodiversity are generally projected to the future, disregarding spatial assessments of current and recent historical climatic change impacts that could actually guide future mitigation and adaptation decisions over more solid basis.

Climate has been changing pervasively since mid 20^th^ century. Global mean surface temperature has increased around 0.85°C in the last 130 years, and precipitation has generally increased in latitudes beyond 30°, but decreased in the tropics since the 1970s [6]. Patterns of climate change are dynamic and highly heterogeneous across the planet, where differential warming or cooling rates occur, thus uniform biodiversity responses across the planet should not be expected [7,8].

One of the main inputs for regional or global climate change assessments are interpolated climatic surfaces at high spatial resolution [9]. High-resolution climatic surfaces have been useful for assessments of biodiversity responses to climate change [10]. Currently, there are some such global databases freely available such as *WorldClim* and *Climond* [9,11]. However, such databases present some shortcomings that impede evaluating the historic trends of climate change and their impacts on biodiversity; for example, they span over a single period of time in the late 20^th^ and early 21^th^ centuries (1950-2000) and their quality and reliability in many regions is compromised because they have been produced with information available in open sources. Nonetheless, many countries or regions hold climatic data unavailable for global analysis. For instance, Mexico has a diverse set of climatic surfaces developed specifically for this country [12,13,14], but although these surfaces improve global datasets as they were built using information from a higher density of weather stations, none of them is useful to evaluate the spatio-temporal dynamics of climate change for being a single average of climatic variables over variable temporal spans [15,16].

In turn, climatic surfaces from different periods of the recent past allow analyses to understand where, when and the magnitude at which climate is changing, and therefore evaluate the implications for biodiversity [17,18,19,20,21,22,23,24,25]. For instance, combining this information with species inventories can be useful to detect drivers of change in biological communities [25,26], and responses of biodiversity to climate change [27]. Historic climate surface can indicate which areas have been stressed by past climate change and thus need conservation action. Also, coupling this information with future projections reduces uncertainty in climatic scenarios, with important implications for science and decision-making [28,29,30]. For example, the analysis of historical climate in Australia permitted to detect recent range shifts in bird communities, giving a better understanding of future biotic responses [31].

Until now, most studies evaluating climate change impacts on biodiversity have focused on future projections of individual species, whereas analyses at other biological levels have been relegated [32]. Therefore, there is a need to analyse climate change impacts at higher biological diversity levels, such as biogeographic provinces, which are natural spatial units that integrate physiographic, evolutionary and ecological features of biodiversity [33]. There is no information available of the impacts of current climate change at this organization level to assess their vulnerability [28]. Therefore, spatial analyses at this scale may provide a range of climatic trends at a regional scale that can serve to establish conservation priorities under current and future climate change [32].

Here, we have developed historical climatic surfaces for three periods that cover most of the 20^th^ and early 21^th^ centuries: *t*_*1*_-1940 (1910-1949), *t*_*2*_-1970 (1950-1979) and *t*_*3*_-2000 (1980-2009) and used these surfaces to describe historical trends of climate change within the 19 biogeographic provinces of Mexico.

## Materials and Methods

### Climate data

We analysed monthly minimum and maximum temperature and accumulated rainfall data gathered from weather stations of the National Meteorological Office, prearranged by Cuervo-Robayo [14], to derive monthly mean climate surfaces for three periods 1910-1949 (*t*_*1*_-1940), 1950-1979 (*t*_*2*_-1970) and 1980-2009 (*t*_*3*_-2000). We selected these periods based on global [34,35] and regional [16] climate change analysis and we also considered the number of stations available for each period [36]. For the first period (*t*_*1*_-1940) we used a 40-year period due to the limited number of available stations. Climate data were averaged and organized into their corresponding periods with the Structuration tool of the Integrated Water Management extension, implemented in the software Idrisi Selva and that is freely available at http://Idrisi.uaemex.mx [37].

We used the ANUSPLIN software version 4.37 [38] to generate countrywide climatic continuous surfaces. This program uses the thin-plate smoothing splines interpolation technique, which integrates climatic and topographic data for making spatial climatic estimations, thus performing better than other interpolation methods [39]. Interpolation estimations are normally poor towards the edges of a region; therefore, the analysis should expand beyond the limits to the target region. Consequently, we included weather stations from southern portions of the United States, northern Belize and Guatemala. Finally, we only used climate data for stations that operated for more than 10 years in at least one variable (temperature and precipitation). We used a second order spline with three independent variables (latitude, longitude, and elevation in km), and a square root transformation for precipitation [14]. The square root transformation reduces positive skewed values and ignore all negative values in precipitation data [38,40]. It also, applies more smoothing to large precipitation values and less smoothing to small precipitation data values [41].

ANUSPLIN produces a list of the stations with largest data residuals that denotes fitting errors. With this list Cuervo-Robayo et al. [14], correct the geographic position for a hundred of erroneous stations by using online gazetteers and Google Earth, however for the last two periods we eliminated about 20 stations that maintained high residuals values during the fitting process. We assessed the accuracy of the fitted surfaces by examining ANUSPLIN diagnostic measures [38]. The signal indicates the degrees of freedom associated with the surfaces, which reflects the complexity of the surface and varies between a small positive integer and the number of stations used to generate the surface [40,42]. Hutchinson & Gessler (1994) suggested that the signal should not be greater than about half the number of data points. Models with a signal below this threshold tend to be more robust and reliable in regions where data are scarce [43]. We also examined the root mean square error (RTMSE) and the square root of the GCV (RTGCV). RTMSE is an optimistic assessment of predictive error since some of interpolation error is removed. On the other hand, it does include error due to the data including quite a few short period means. The true error is somewhere between the RTMSE and the RTGCV error [38]. Gridded climate surfaces were generated with the function LAPGRD, using a 30-arc seconds of spatial resolution from the GTOPO30 digital elevation model (https://lta.cr.usgs.gov/GTOPO30). As an additional output, we derived 19 bioclimatic variables for each period with the dismo library [44]of R software [45]. These variables include annual, quarterly, and monthly summaries of temperature and precipitation that represent more biologically meaningful variables than the original climate surfaces and have been widely used in several studies of climate change impacts on species and ecosystems [9].

For the first period (*t*_*1*_-1940), we used 803 stations for precipitation and 500 for minimum and maximum temperatures. In the second period (*t*_*2*_-1970), the number of stations for precipitation and temperatures were 3411 and 3670, respectively. For the last period (*t*_*3*_-2000), the number of stations increased to 3870 and to 4200 for precipitation and temperature, respectively. Data used for the analysis are available upon request, and the original dataset is available at the National Meteorological Office (http://smn.cna.gob.mx/es/).

In order to evaluate differences between periods we firstly *z*-standardized monthly precipitation, maximum and minimum temperatures grids. Standardized monthly grids were than used to assess differences between periods using a Discriminate Analysis Function [46]. Additionally, to evaluate if differences in climate between *t*_*2*_-1970 and *t*_*3*_-2000 were caused by differences in the number of weather stations used for each period, we developed a set of climate surfaces in which we used the same number of weather stations for both periods. With these evaluations, we found that differences were not due to the number of stations, thus we included more stations for the third period, which improved model signal [14,47]. We did not compare our climate surfaces with other regional or global models, because interpolation techniques, temporal coverage, number and climate stations differ and can therefore lead to non-objective comparisons.

### Biogeographic provinces of Mexico

Mexico is recognized as one of the top-10 megadiverse countries in the world [48]; this extraordinary diversity is due to its high environmental heterogeneity and the confluence of three main biotic components; the Neartic and Neotropical regions, and the transition zone. Biotic components rarely have a “unique” origin, but are usually sets of elements of different affinities, which have been integrated into the course of biotic evolution [49]. We characterize climate change in the biogeographic provinces because each region is dominated by biotic components with distinctive evolutionary origins that could respond to climate change based on their affinity. The Nearctic region encompasses the arid subtropical areas of the north, which is assigned to the provinces of California, Baja California, Sonora, Altiplano Mexicano and Tamaulipeca. The Neotropical region contains the humid and sub-humid tropical areas of the south, assigned to the provinces of Costa del Pacífico, Golfo de México, Oaxaca, Altos de Chiapas and the Península de Yucatán, which includes Yucatán and Petén. The Transition zone includes the mountainous area in the central portion of the country: Sierra Madre Oriental, Sierra Madre Occidental, Eje Volcánico Transmexicano, Cuenca del Balsas and Sierra Madre del Sur [49,50].

A map of the biogeographic provinces of Mexico (Fig 1) was obtained from the *Comisión Nacional para el Conocimiento y Uso de la Biodiversidad* (CONABIO 1997 http://www.conabio.gob.mx/informacion/gis/). This regionalization was based on the distributional pattern of four taxonomic groups (vascular plants, amphibians, reptiles, and mammals) and the main morpho-tectonic features of Mexico [51]. Each unit represents a relatively homogeneous area that concentrates high levels of endemic species sharing similar historical, physiographic, climatic, edaphic, and vegetation features [50].

**Fig 1.**
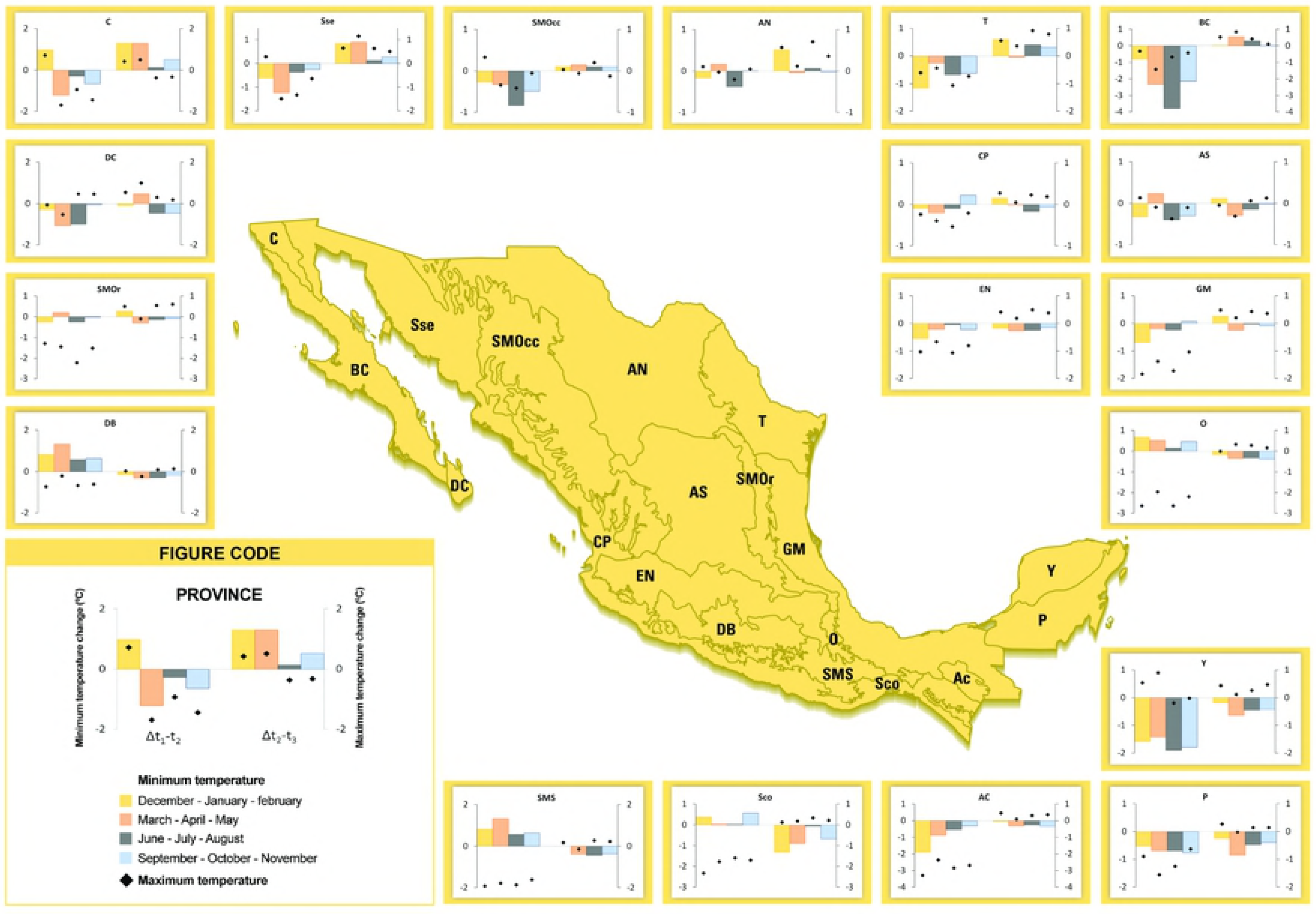
Seasonal maximum (♦) and minimum (bars) temperature change from *t*_*1*_-1940 to *t*_*2*_-1970 (Δ*t*_*1*_-*t*_*2*_) and from *t*_*2*_-1970 to *t*_*3*_-2000 (Δ*t*_*2*_-*t*_*3*_) within the biogeographic provinces of Mexico. DB: Depresión del balsas, SMOr: Sierra Madre Oriental, DC: Del Cabo, C: California, Sse: Sonorense, SMOcc: Sierra Madre Occidental; AN: Altiplano Norte, T: Tamaulipeca, BC: Baja California, CP: Costa del Pacífico, AS: Altiplano Sur, EN: Eje Neovolcánico, GM: Golfo de México, O: Oaxaca, Y: Yucatán, P: Petén, AC: Altos de Chiapas, Ssc: Soconusco and SMS: Sierra Madre del Sur. Negative values indicate a decrease in temperature from the previous period and positive values indicate an increase.

### Climate change rates and trends

To characterize seasonal change rates from *t*_1_1940 to *t*_*2*_-1970 and from *t*_*2*_-1970 to *t*_*3*_-2000, we estimated the mean monthly climatic profile values within each biogeographic province with the software Earth Trends Modeler (ETM) of Idrisi Selva (Eastman 2012). These values corresponded to the mean value of the climate variable over each province. Monthly profiles were averaged in four seasons: (i) December, January, February (DJF), (ii) March, April, May (MAM), (iii) June, July, August (JJA), and (iv) September, October, November (SON). With these profiles, we estimated the difference between periods for each season as follows (using as an example maximum temperature - *maxT-* notation):

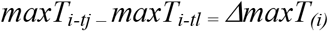

where _*i*_ represents the season of _*tj*_ and _*tl*_ periods. Differences were estimated for precipitation (*PPT*), maximum temperature (*maxT*) and minimum temperature (*minT*). Differences in precipitation were estimated in millimetres and percentage (S1 eq. 1), and differences in temperature in °C.

We assessed spatial trends of monthly temperature and precipitation during these three periods, continuously from January of the *t*_*1*_-1940 to December of the *t*_*3*_-2000, with a Mann-Kendall test. This test is a non-linear tendency indicator that measures the degree at which a trend is consistently increasing or decreasing. The Mann-Kendall statistic is simply the relative frequency of increases minus the relative frequency of decreases [52], ranging from - 1 (always decreasing) to +1 (always increasing) and a value of 0 indicates no consistent trend. This test is useful to detect areas that have been exposed to constant climatic changes. We calculated Mann Kendall test for each pixel of the z-standardized monthly precipitation, maximum and minimum temperatures grids in the Series Trend Analysis tab of ETM trends. None significant (P < 0.05) pixels were excluded in order to have a raster with significant trends only. However; even when the Mann Kendall test can be used to measure the consistency of exposure from a given location to climate change, it does not represent the magnitude of change.

## Results

### Climate surfaces

ANUSPLIN diagnostic measures indicated an overall adequate fit of spline models for the three climatic periods. The average ratio of the signal to the number of data points was <0.5 for monthly temperatures and precipitation (S1). For the first period, a small number of climate stations (<900) with precipitation data were available, so we only eliminated the stations with higher residual error to maintain a reasonable number of stations (>800). The average signal for precipitation is above the permitted threshold indicating that this variable is too complex to be adequately represented by the data (Hutchinson et al. 2009), but for some particular months (i.e., from January to May) the ratio signal was acceptable (S1). Therefore, period *t*_*1*_-1940 must be used with caution. Overall, the monthly average RTMSE for both temperatures were below 0.6 and below 10 mm for precipitation.

We found differences in the standardized grids of precipitation (Wilks’ Lambda = 0. 295, F_*df* = 24,178478_ = 6249.079, *P* < 0.001), maximum (Wilks’ Lambda = 0.419, F_*df* = 24,178478_ = 4050.582, *P* < 0.001) and minimum temperature (Wilks’ Lambda = 0.398, F_*df* = 24,178478_ = 4346.174, *P* < 0.001) between the time periods. Monthly climate surfaces and bioclimatic variables for each period are freely available at: http://www.conabio.gob.mx/informacion/gis/, using the search tab and the words: *“clima histdrico*” or “historic climate”.

### Climate change rates in biogeographic provinces

We estimated climate change rates and spatial trends for the 19 biogeographic provinces of Mexico. Climate change rates represent average value of change in biogeographic provinces, whereas spatial trends of change refer to the geographically consistent tendency of increase or decrease in the variables within each pixel or region, thus it does not indicate the magnitude of change.

Across Mexico, maximum and minimum temperatures have increased since *t*_*2*_-1970 in most provinces and for almost all seasons, even when at the beginning of the century maximum temperature decreased mainly in the southern provinces of the country (Fig 1). Within seasons the pattern is similar to the general trend, we observed a notable increase of temperatures throughout the evaluated periods, particularly for the second half of the 20^th^ century, although changes in minimum temperature within seasons were not consistently showing warming. This happened mainly in the biogeographic provinces of Depresión del Balsas, Oaxaca, Sierra Madre del Sur, Oaxaca and Socunusco. Moreover, the first two seasons-corresponding to winter and spring (DJF and MAM) - warmed up more drastically in the northern provinces in the last 30 years than in the rest of the country (Fig 1).

In general, precipitation rate of change has been positive throughout the century. However, on average, precipitation rate slowed down in the transition *t*_*2*_-1970-*t*_*3*_-2000 compared to *t*_*1*_-1940-*t*_*2*_-1970. In addition, in several southern biogeographic provinces, precipitation rate was negative in *t*_*2*_-1970-*t*_*3*_-2000, mainly during the spring and autumn (MAM; SON; Fig 2). Mean precipitation rate was higher from *t*_*1*_-1940 to *t*_*2*_-1970 in five biogeographic provinces (i.e., Sierra Madre del Sur, Costa del Pacífico, Oaxaca, Soconusco and Golfo de México) compared to the mean precipitation rate during *t*_*2*_-1970-*t*_*3*_-2000. In the remaining provinces, precipitation rate was positive in *t*_*1*_-1940-*t*_*2*_-1970 but with a lower rate compared to the transition *t*_*1*_-1940-*t*_*2*_-1970 (Fig 2).

**Fig 2.**
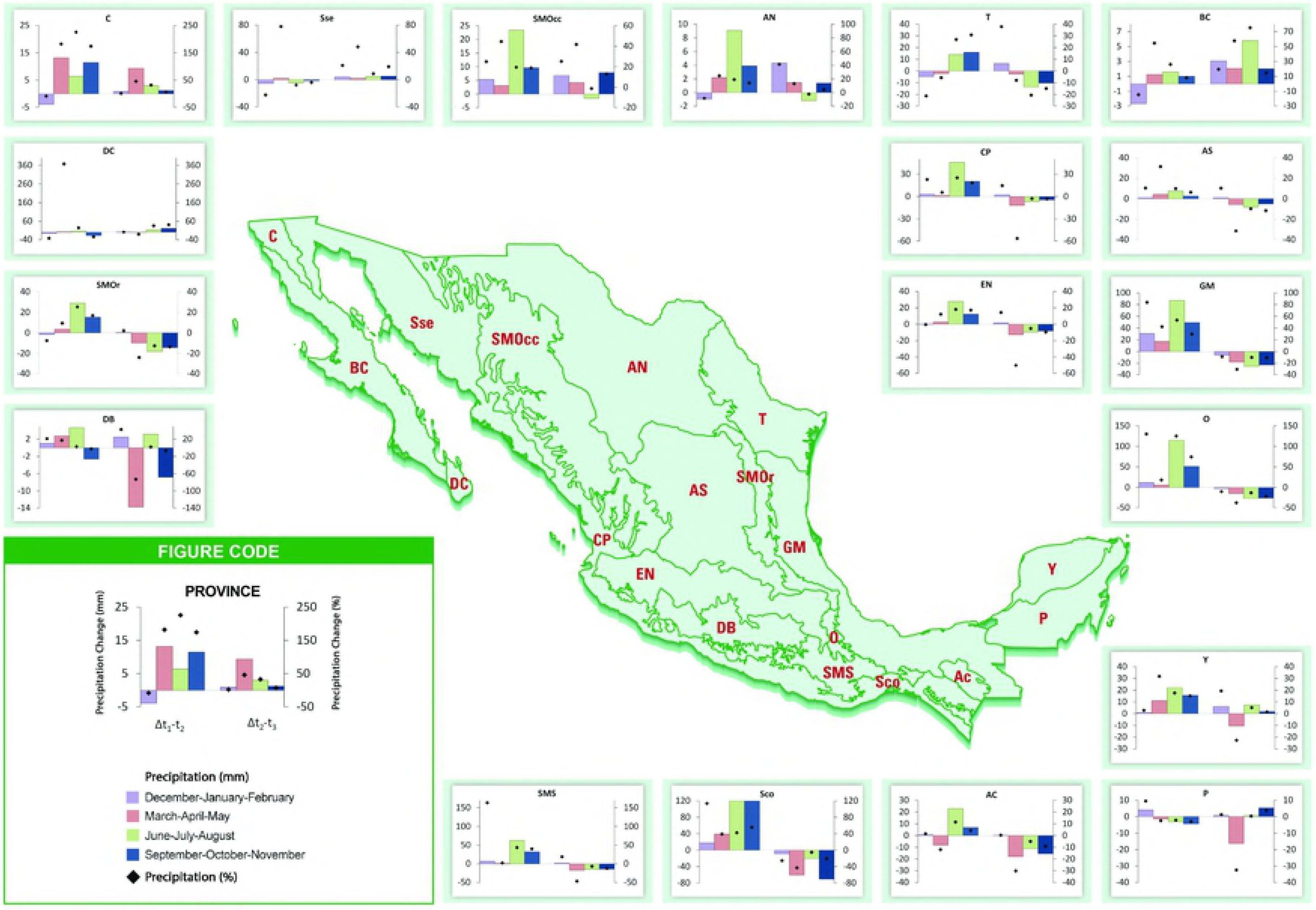
Seasonal precipitation in % (♦) and in milimeters (bars) change from *t*_*1*_-l940 to *t*_*2*_-1970 (Δ*t*_*1*_-*t*_*2*_) and from> *t*_*2*_-1970 to *t*_*3*_-2000 (Δ*t*_*1*_-*t*_*3*_) within the biogeographic provinces of México. DB: Depresión del balsas, SMOr: Sierra Madre Oriental, DC: Del Cabo, C: California, Sse: Sonorense, SMOcc: Sierra Madre Occidental; AN: Altiplano Norte, T: Tamaulipeca, BC: Baja California, CP: Costa del Pacífico, AS: Altiplano Sur, EN: Eje Neovolcánico, GM: Golfo de México, O: Oaxaca, Y: Yucatán, P: Petén, AC: Altos de Chiapas, Ssc: Soconusco and SMS: Sierra Madre del Sur. Negative values indicate a decrease in temperature from the previous period and positive values indicate an increase.

### Mann-Kendall trend analysis

The Mann-Kendall analysis showed significant trends in over 50% of the climatic variables evaluated (Fig 3), but these trends were uneven. Strong trends of Mann-Kendall values (>0.5 and <−0.5) were at a low proportion. These happened mainly in the Altiplano del Norte, Sierra Madre Occidental, Sonorense, Baja California, Peten for precipitation plus Yucatán, Sierra Madre del Sur and Eje Neovolcánico for both temperatures.

**Fig 3.**
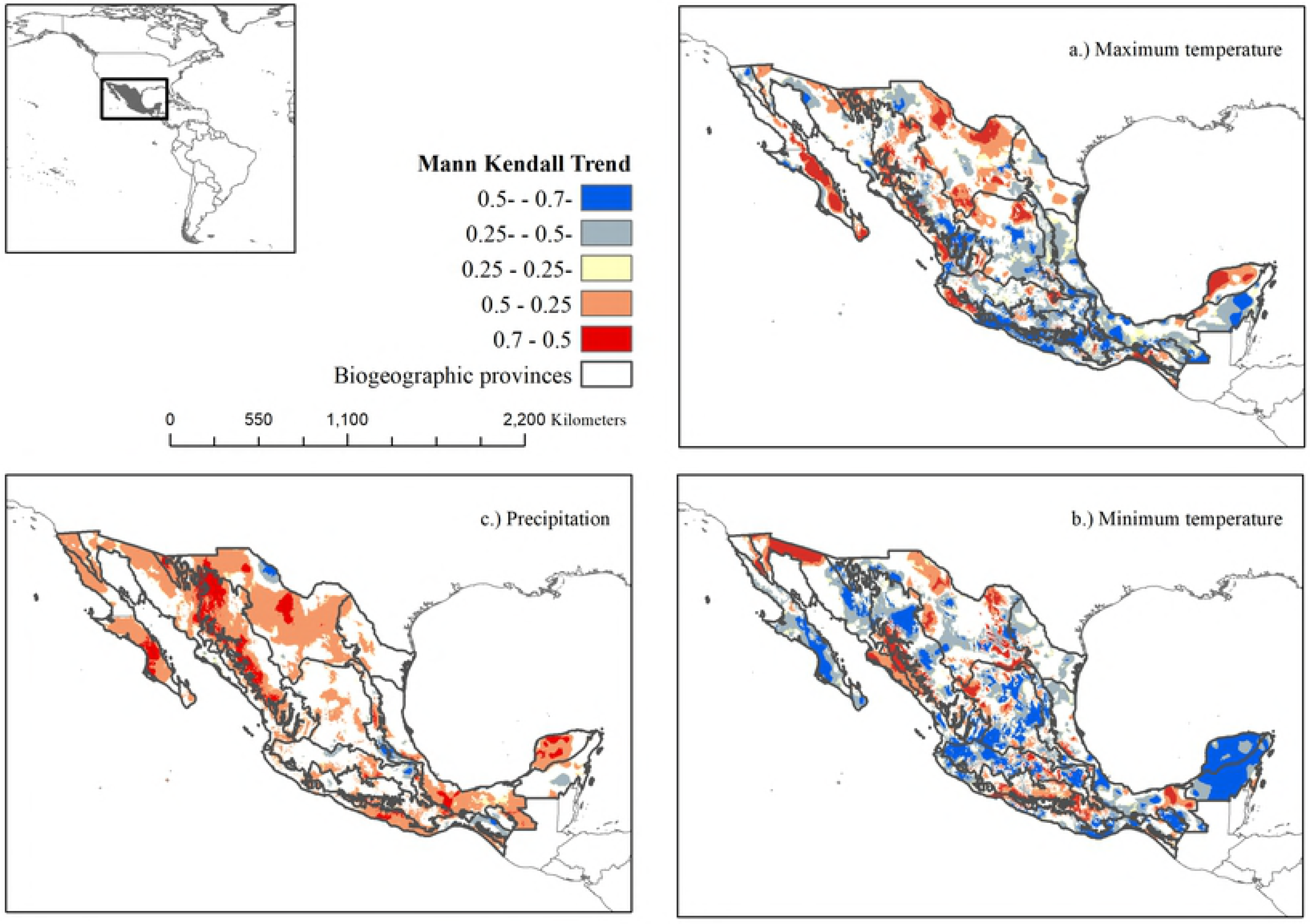
Mann-Kendall trend analysis for three climate variables in Mexico between *t*_*2*_-1910 to *t*_*2*_-1970 and *t*_*2*_-1970 to *t*_*3*_-2000. In parenthesis, the percentage of area occupied by each category of trend. Values above 0.1 and under ^-^0.1 represent positive and negative trends, respectively. Values within 0.1-^-^0.1, represent no consistent trend.

Positive trends for the minimum temperature where particularly consistent in the mountainous regions of the country, in the highest peaks of the Eje Neovolcánico province (i.e., Nevado de Toluca, Popocatepetl and Iztaccíhuatl volcanoes). On the other hand, the negative trends observed for the minimum temperature occurred mainly in the same areas where a negative rate of change were also observed (Fig 1), namely, the Yucatán, Los Altos de Chiapas and the Petén provinces.

Finally, we identified a consistent warming up trend in the maximum temperature in the California, Baja California, Sonorense, Del Cabo, and Tamaulipeca provinces throughout the century. In the transition *t*_*1*_-1940-*t*_*2*_-1970 we observed a warming trend in the northern provinces extending 63% of the country (Fig 3) and a cooling trend in southwestern Mexico covering around 11% of the country. Finally, we identified a warming trend in the highest elevations of the Eje Neovolcánico and Golfo de México, close to Taumaulipeca province.

## Discussion

Climate change is currently the environmental topic of greatest concern worldwide, thus extensive research efforts are underway. Surprisingly, formal evaluations of historical climate change have been seldom carried out at a regional level [18]. To accomplish this goal, historical climatic surfaces have proven useful to analyse climatic variation over the century and its effects on biodiversity and agrobiodiversity at all biological levels [41].

We developed historic climatic surfaces to evaluate the extent at which Mexican biogeographic provinces have been exposed to climate change during the 20^th^ and early 21^st^ centuries. We identified a general positive trend of change in precipitation throughout the century, although the rate decreased since *t*_*2*_-1970. Countrywide, Mexico has been projected to maintain medium to high climatic stability [32], but, as expected, rates of change and trends have not been historically uniform [53]. Our results indicate that for the three mayor biogeographic regions of the country Nearctic, Neotropical and Transition; northern provinces at the Nearctic region are the ones that have been more exposed to climate change due to faster rates of change and consistent positive trends. These could lead to large changes in species compositions and biodiversity [1], as it has been observed for endemic birds of Mexico [25].

The Transition region includes the Sierra Madre Occidental, Sierra Madre Oriental and the Eje Neovolcánico provinces, which are characterized by an admixture of species of Nearctic and Neotropical origin [49]. The provinces within this region harbours diverse types of temperate vegetation, such as oak, pine and tropical cloud forests; all of them, but particularly the latter, highly vulnerable to climate change [54,55]. Thus, the climatic changes detected in this study are important to establish priorities for conservation.

Our analyses demonstrate that the Neotropical region (Costa del Pacífico, Golfo de México, Oaxaca, Sierra Madre de Sur, and Soconusco provinces) has been exposed to a more pronounced decline in precipitation rates, probably as a consequence of an increase in the frequency and intensity of El Niño events in the last two decades [56]. Conversely, precipitation and temperature [16] have shown a positive trend in some provinces of the north, namely, California, Sonorense, Altliplano del Norte, and Tamaulipeca. In these regions precipitation is counterbalanced by evapotranspiration, and the combined effect of an increase in precipitation and temperature can causes a higher vapour pressure deficit and evaporation, reducing water availability [53], as has been observed in Tamaulipeca, Baja California and Sonorense provinces [57,58]. Arid and semiarid provinces are highly dependent on water availability, regulating net ecosystem productivity [53,59] and agriculture [32,57]. Human population and agriculture have increased in these provinces, exceeding water demand in some areas [56].

Our results confirm that Mexico has generally warmed up during at the end of the 20^th^ and beginning of the 21^st^ centuries (*t*_*2*_-1970 and *t*_*3*_-2000), as observed by Pavia *et al.* [16] and Englehart and Douglas [15]. However, these authors found that warming has not been consistent throughout the country during the 20^th^ century, as we also observed. They reported that cooling occurred mainly in the central and southern provinces of the country at mid century (1940-1969). In turn, we observed a decrease of the maximum temperature at the Sierra Madre Oriental and Soconusco provinces from *t*_*1*_-1940 to *t*_*2*_-1970, but a clear increase since *t*_*2*_-1970 that was also observed in the Sierra Madre Oriental, Golfo de México, Altos de Chiapas and Yucatán provinces. The Sierra Madre Oriental harbours the richest coniferous forests in the world [60], and is projected to be affected by climate change [61], thus priority attention in this region is advised [60].

There was no consistent trend in most of the Altiplano del Sur and Eje Neovolcánico provinces, although within the latter we identified a positive trend in all volcanic areas. The rapid retreat of glaciers during the 20^th^ century confirms this observation [62]. The confluence of flora and fauna from the Nearctic and Neotropical provinces makes this province particularly rich [50]. Many species within it hold naturally narrow distributional ranges, and coupled with the fact that this province concentrates the highest human density and largest urban nuclei of the country, its biodiversity is highly vulnerable to the synergistic effects of multiple stressors, including climate change [7]. Furthermore, many species’ survival will depend on their capacity to keep pace with climate, however, their limited ability to respond to climatic changes via elevation range shifts due to human-induced obstacles make them highly vulnerable to the current warming event [32,53].

Understanding recent climate and ecological changes can be useful to focus research efforts and conservation actions. Changes in climate are now occurring simultaneously with other environmental disruptions. It will not be possible to fully understand biodiversity responses to climate change without taking into account the interactions with other components of environmental change [63]. Species responses will depend on their exposure and sensitivity to other human-induced pressures, their inherent capacity to adapt to new conditions, the magnitude and speed of environmental changes, and time lags in their responses [53].

Our aim is that this contribution serves as a baseline for improving our knowledge regarding climate change in Mexico through time in a spatially explicit fashion, to identify specific areas where climate change is occurring, its direction and magnitude. In summary, climate change is occurring unevenly in Mexico, being the provinces at the Nearctic region more affected by a higher warming rate and a sustained trend compared to provinces at the Neotropical and Transition regions. In turn, precipitation has increased across the country, although this rate has slowed down towards the end of the 20^th^ century, particularly in the provinces of the Neotropical region. Finally, this information can be used to improve regional projections of future climate trajectories, which would provide scientists and authorities with more reliable data and information for making better decisions in the face of climate change.

## Caveats

We have generated statistically robust historical climatic surfaces for Mexico, which are considered acceptable for this type of model [38] and comparable to two other climatic surfaces developed for Mexico for different time frames [12,14]. However, *t*_*1*_-1940 was interpolated with a reduced number of weather stations presenting larger inconsistencies than the other two periods [14]. Therefore, we advise caution in the use of this particular dataset. Reliability of products involving historical data, as the ones presented here, is limited by deficiencies and inaccuracies of data itself [41].

## Acknowledgments

The Mexican Consejo Nacional de Ciencia y Tecnología (CONACyT) provided a Ph.D. scholarship to A.P.C-R (No. 217650) and DGAPA-UNAM a Postdoctoral scholarship to C.U. This work was funded by PAPIIT-DGAPA-UNAM (Grant number IN-225010). We thank Crystian Venegas for helpful comments on the methodology. The Servicio Meteorológico Nacional (SMN) provided data from the weather stations. We also thank the Comisión Nacional para el Conocimiento y Uso de la Biodiversidad (CONABIO) for being the repository of the climate surfaces developed in this study.

## Author Contributions

**Conceptualization**:

**Data curation: ACR, MGA and AMM**.
**Formal analysis: ACR and EMM**
**Funding acquisitions: ACR, OTV and EMM**.
**Investigation: ACR, CU, MGA, OTV and EMM**.
**Methodology: ACR, MGA, AMM and EMM**.
**Project administration: ACR and EMM**
**Resources: ACR, OTV, EMM**
**Software: OTV**
**Supervision: OTV and EMM**.
**Validation: ACR, MGA and EMM**.
**Visualization: ACR**
**Writing-original draft: ACR, CU and EMM**.
**Writing-review & editing: ACR, CU and EMM**.

## Supporting information

### S1. Equation and ANUSPLIN statistics

This file contains the formula used to estimate the precipitation and two tables.

